# Aggregating gut: on the link between neurodegeneration and bacterial functional amyloids

**DOI:** 10.1101/2024.11.26.624671

**Authors:** Alicja W. Wojciechowska, Jakub W. Wojciechowski, Kinga Zielinska, Johannes Soeding, Tomasz Kosciolek, Malgorzata Kotulska

## Abstract

Amyloids are insoluble protein aggregates with a cross-beta structure, which are traditionally associated with neurodegeneration. Similar structures, named functional amyloids, expressed mostly by microorganisms, play important physiological roles, e.g. bacterial biofilm stabilization. Using a bioinformatics approach, we identify gut microbiome functional amyloids and analyze their potential impact on human health via the gut-brain axis. The results point to taxonomically diverse sources of functional amyloids and their frequent presence in the extracellular space. The retrieved interactions between gut microbiome functional amyloids and human proteins indicate their potential to trigger inflammation, affect transport and signaling processes. We also find a greater relative abundance of bacterial functional amyloids in patients diagnosed with Parkinson’s disease and specifically a higher content of the curli amyloid protein, CsgA, in Alzheimer’s disease patients than in healthy controls. Our results provide a rationale for the tentative link between neurodegeneration and gut bacterial functional amyloids.

## Introduction

The influence of the human gut microbiome on health, including the occurrence of neurodegenerative disorders, has been well established by now, Fang et al. (2020) Gilbert et al. (2018), Knight et al. (2017). The microbiome is responsible for the supply of multiple substances, such as folate, riboflavin (vitamin B2), vitamin B12 or short-chain fatty acids, like butyrate, which have anti-inflammatory properties and serve as an energy source for the host, LeBlanc et al. (2013), Fusco et al. (2023). Furthermore, it shapes immunological responses by modulating cytokine production and lymphocyte activation, Schirmer et al. (2016), Ivanov et al. (2009).

Recent studies showed that many bacterial strains are capable of producing proteins that are known as bacterial functional amyloids, Otzen and Riek (2019). Their structures are strikingly similar to human misfolded proteins forming plaques in the brains of patients suffering from neurodegenerative diseases. However, unlike human pathological amyloids, the bacterial functional amyloids support the bacteria by performing vital functions, including biofilm formation, adhesion or signaling, Levkovich et al. (2020).

It has been proposed that bacterial functional amyloids could indirectly affect the onset and progression of neurodegenerative diseases. This mechanism has been proposed in Parkinson’s disease (PD), where interactions between bacterial functional amyloids and a human protein alpha-synuclein, whose aggregation contributes to the disease onset, increased the level of disease incidence, Friedland and Chapman (2017). Due to the structural similarity of human misfolded proteins and bacterial functional amyloids, they are capable of affecting the aggregation rates of their pathological human counterparts, Burdukiewicz et al. (2023), Konstantoulea et al. (2021). Bacterial functional amyloids may trigger misfolding of human amyloids and their aggregation, Elkins et al. (2024). In this case, pathological aggregation could start in the enteric nervous system and then propagate through the vagus nerve to the brain, Braak et al. (2003). Several studies support this hypothesis; alpha-synuclein aggregates appear early in the PD course in the enteric nervous system and they are correlated with disease severity, suggesting that disease may start in the gut, Lebouvier et al. (2010), Shannon et al. (2011). Moreover, application of inhibitors of the curli CsgA protein (gut bacterial functional amyloid, produced e.g. by *Escherichia coli*) reduces aggregation of alpha-synuclein in the mouse brain model, Sampson et al. (2020). Furthermore, biofilm-related bacterial functional amyloids are more abundant in PD patients than in healthy controls. They can also co-localize with alpha-synuclein in neurons and increase its aggregation, Fernández-Calvet et al. (2024).

The potential role of bacterial functional amyloids in neurodegeneration is, however, not limited to triggering aggregation. Neurodegenerative diseases are characterized by gut dysbiosis, dysfunction and inflammation, Kelly et al. (2013), Heravi et al. (2023), Hirayama and Ohno (2021). Bacterial functional amyloids, due to their structural similarity to human pathological amyloids, could have similar cytotoxic activity in humans. Moreover, their presence, especially during dysbiosis, could be another pro-inflammatory factor leading to abnormal gut permeability, Bucianntini et al. (2002).

Although many studies have shed light on the potential effects of bacterial functional amyloids on neurodegeneration, multiple questions remain unaddressed. Are bacterial functional amyloids present on a large scale in the human gut and, if so, which bacteria produce them? Are they more abundant in microbiomes of patients with neurodegenerative diseases than in controls? And, finally, which molecular pathways could be affected by gut bacterial functional amyloids? Answers to these questions could broaden our understanding of the onset of neurodegeneration and related molecular mechanisms governing the role of the microbiome, potentially enabling the development of novel therapeutic strategies or biomarkers for early diagnosis.

In this work, we aim to better understand the potential influence of bacterial functional amyloids and their link to neurodegeneration. Through a detailed computational analysis of available datasets, we develop an atlas of predicted bacterial functional amyloids expressed by the human microbiome. Using the atlas, we investigate the abundance of bacterial functional amyloids in the microbiome metagenomic samples of patients with Parkinson’s disease (the most studied in the context of the potential influence of microbiome), Alzheimer’s disease, and bacterial brain infection Cryptococcal Meningitis (negative control). Finally, we predict human interactors of gut bacterial functional amyloids and reveal that they can affect human cell signaling, transport, and even immunity.

## Results

### A vast array of bacterial functional amyloids may be present in the human gut microbiome

To identify putative bacterial functional amyloids in the human gut, we relied on the fact that their aggregation propensity should be evolutionarily conserved, Otoo et al. (2008), Dueholm et al. (2013), Dueholm et al. (2012). We performed the identification of homologous sequences of known *bacterial functional amyloids* (*BFA* dataset) in the human gut microbiome (*UHGP* dataset) (Fig. 1), Nowakowska et al. (2023), Almeida et al. (2020). The homology search yielded 10,249 sequence matches and 9,541 unique sequences in total. Longer proteins, such as biofilm-related or DNA/RNA-related, provided us with multiple homologous sequences due to different domain matching (WapA: 730, HelD: 714, Aap: 714, YghJ: 714, SuhB: 714, Bap: 713, Smu_63c: 711, PAc: 702, Tuf: 714). For three proteins (alpha phenol-soluble modulins and RdlB) from the *BFA* dataset, we found no homolog in *UHGP*. The mean sequence identity was 45% with a standard deviation of 22%. The mean E-value was 5×10^-6^.

**Fig. 1.**
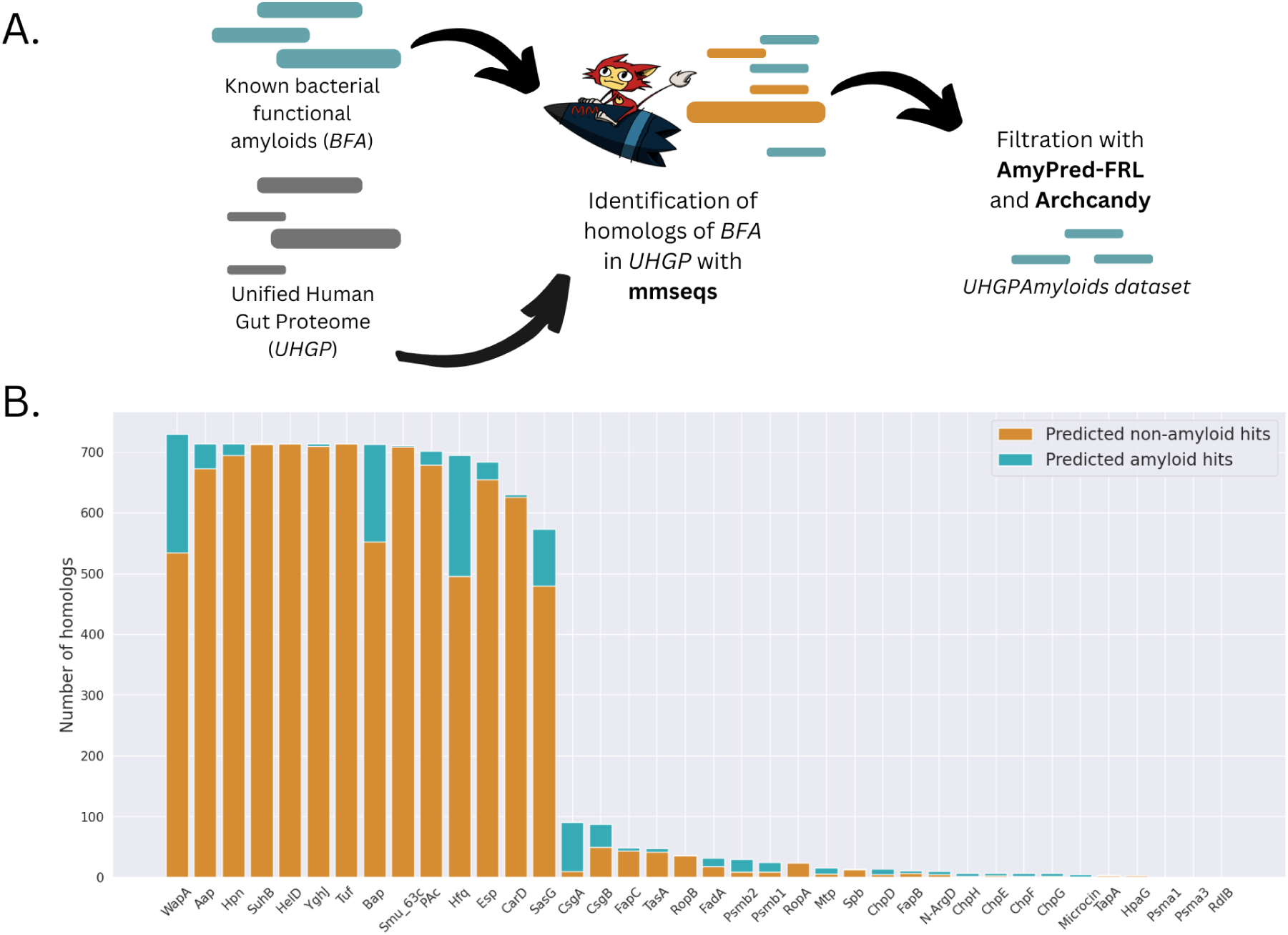
Identification of bacterial functional amyloids in the *UHGP* dataset. A. Pipeline for the search of bacterial functional amyloids in human gut microbiome, B. Number of homologs found per *BFA* protein.

The identified homologs were tested for their aggregation propensity using the full-length sequence predictor AMYPred-FRL, Charoenkwan et al. (2022), which discarded 90% of the sequences, leaving 855 proteins. In general, the percentage of predicted amyloids varied for different proteins. We observed no correlation between AMYPred-FRL score and any of the homology search parameters (Pearson’s |R| < 0.2). We identified beta-arch motifs with ArchCandy, Ahmed et al. (2015), and 50 (6%) sequences in which none was found were discarded. The resulting final dataset of predicted amyloids was labeled as *UHGPAmyloids*. To identify the sequence diversity of *UHGPAmyloids,* we clustered sequences at 90%, 80% and 70% identity, which gave 412, 302 and 243 sequences, respectively. The relatively modest drop in the number of sequences after clustering indicates that although the homology search, in essence, looks for similar proteins, we managed to reach beyond the known world of bacterial functional amyloids.

### Bacterial functional amyloids have diverse taxonomic origin

For each protein from *BFA*, we analyzed the taxonomic origin of all its homologs present in *UHGPAmyloids* and calculated associated Shannon’s entropy (Fig. 2). In addition, we compared the abundance of different phyla between *UHGPAmyloids* and *UHGP* with Fisher’s test (Supplementary Table 1). We observed that although the *UHGP* proteome is mostly produced by Firmicutes, Bacteroidota, Proteobacteria and Actinobacteria (Fig. S1), predicted bacterial functional amyloids are not evenly distributed between these groups (Fig. 2A). Specifically, we observed an enrichment in the following: Firmicutes, Proteobacteria, Fusobacteria. On the other hand, the *UHGPAmyloids* were depleted in: Actinobacteriota and Bacteroidota. No statistical difference was found for Campylobacterota and Myxococcota.

**Fig. 2.**
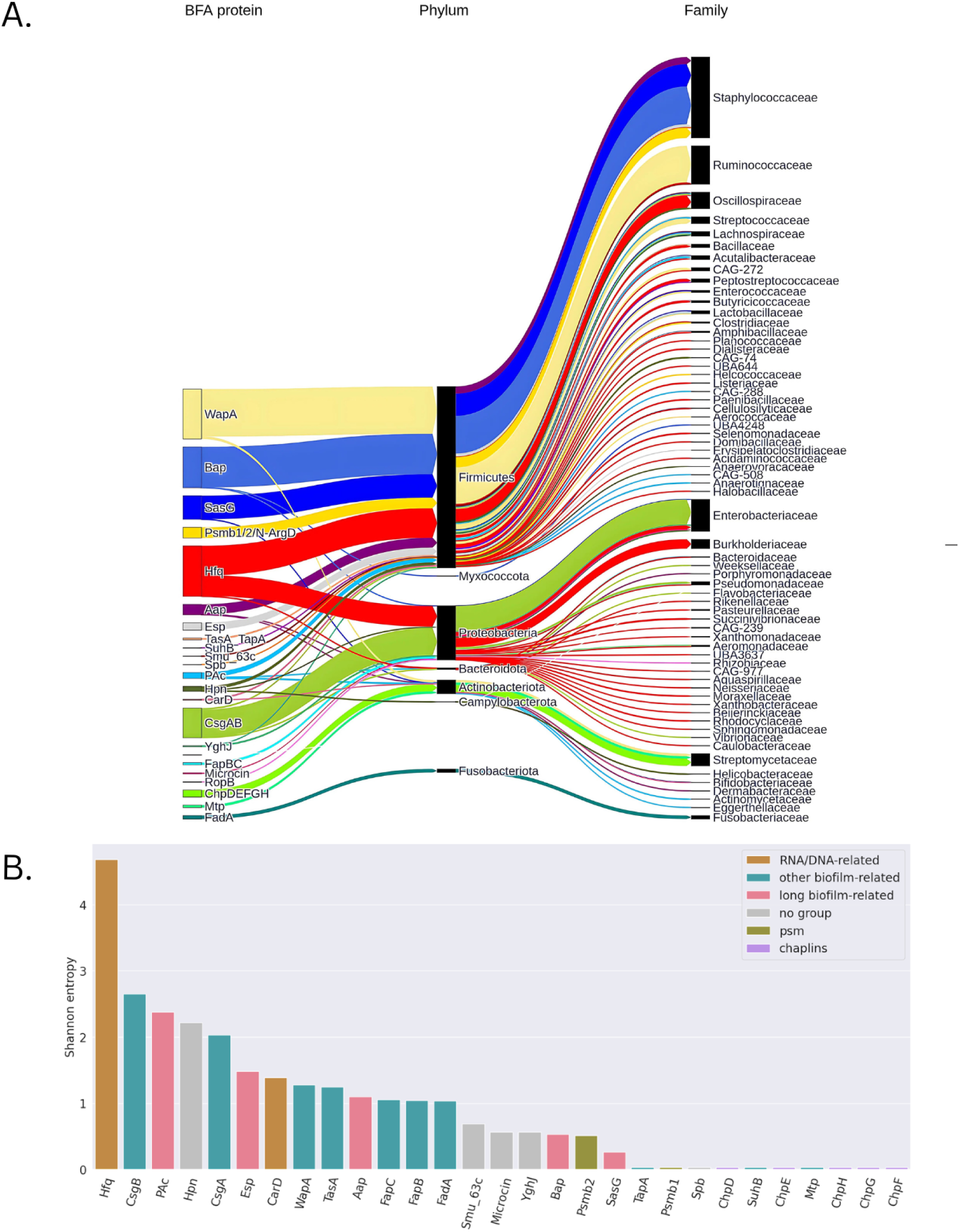
Taxonomic distribution of *UHGPAmyloids*. A. Visualization of taxonomic origin of homologs of *BFA*’s proteins on the phylum and family level. B. Shannon entropy for different groups of homologs.

In the next step, we studied the taxonomic distribution of identified homologs. Some identified bacterial functional amyloids were widely spread across the bacterial tree of life, while others turned out to be specific to certain taxa (Fig. 2A). For example, homologs of chaplins and Mtp were found exclusively in *Streptomycetes*. These microorganisms are capable of producing numerous antibiotics, antiproliferatives, and immunosuppressants, and are hypothesized to have been an important part of the human microbiota in the past, but nowadays are present in low abundance, Bolourian and Mojtahedi (2018). On the other side of the spectrum, the highest taxonomic diversity was found for homologs of the RNA-binding protein Hfq and some biofilm-related proteins, which span across different phyla. However, even in the group of biofilm-related proteins, some variation could be observed. Long biofilm-related proteins WapA, Bap, PAc, Aap and Esp were identified predominantly in Firmicutes with much smaller representation in Proteobacteria, while short biofilm-related proteins, like CsgA and CsgB, were found almost exclusively in Proteobacteria. Interestingly, the changes in the ratios between these two phyla were observed in many human diseases, Shin et al. (2015).

### Gut microbiome of Parkinson’s disease patients is enriched with bacterial functional amyloids

We examined the abundance of identified bacterial functional amyloids (determined in the previous section - *UHGPAmyloids*) in the microbiomes of patients with neurodegenerative (Parkinson’s disease (PD) and Alzheimer’s (AD)) and microbiome-associated inflammatory disease (Cryptococcal meningitis (CM)). We used three metagenomic studies of PD patients and their respective control groups (two datasets collected in different parts of the USA and one in Germany), Bedarf et al. (2017), Wallen et al. (2022), Boktor et al. (2023). To supplement the analysis, we also included one metagenomic study of Alzheimer’s disease (AD) patients and patients suffering from Cryptococcal meningitis (CM), which is an infectious brain disease involving inflammation but not amyloid aggregates deposition, Li et al. 2023. Laske et al. (2022). For all datasets we extracted the original reads, performed quality control, assembled them and searched for homologs of *UHGPAmyloids* (see Methods for details). In all three studies involving PD, we found significantly more bacterial functional amyloids in samples from PD patients compared to the controls (Mann-Whitney U-test p-value < 0.01) (Fig. 3A). In neither the AD nor the CM datasets was there a significant difference in general abundance of amyloid proteins between groups.

**Fig. 3.**
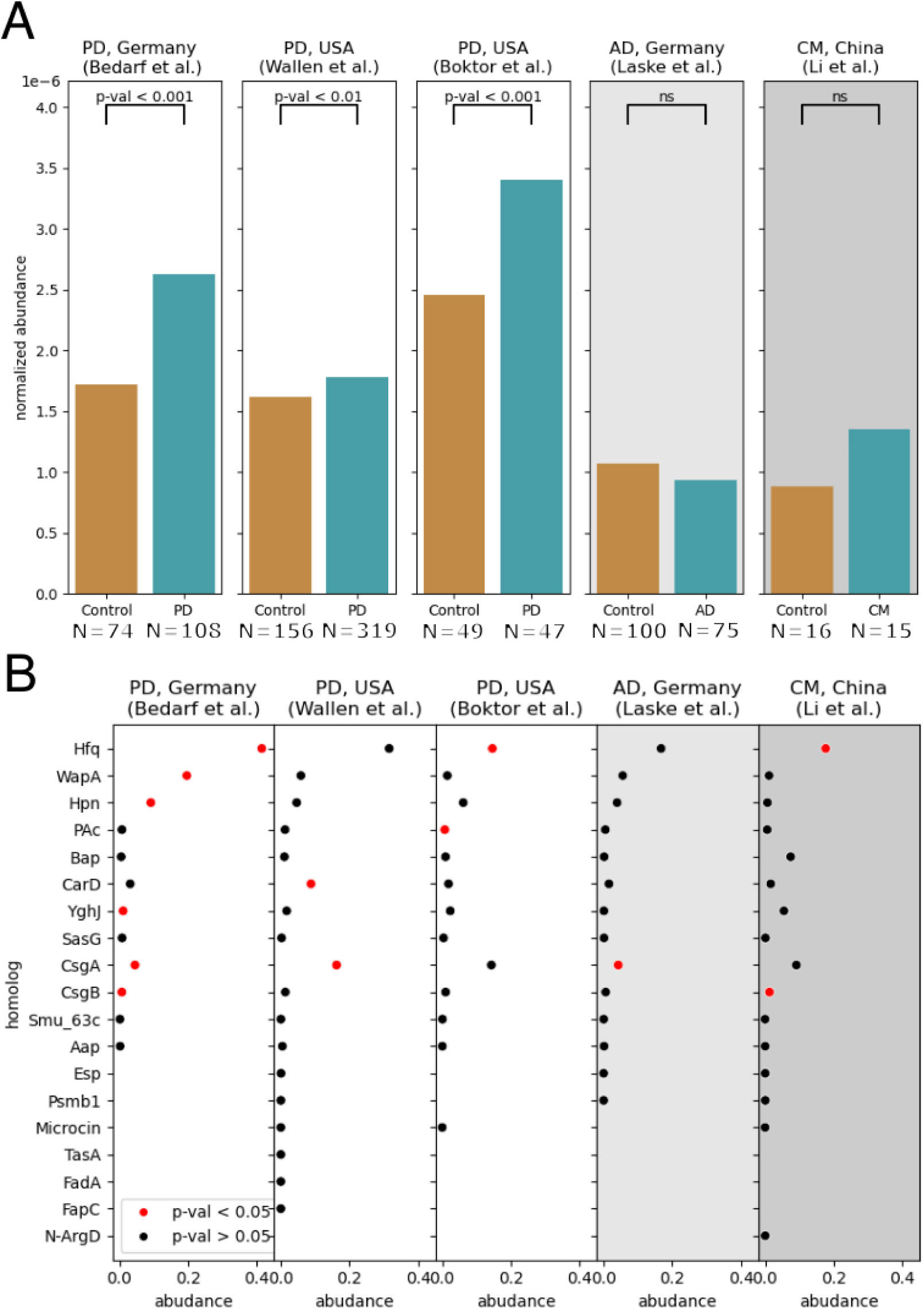
Abundance of bacterial functional amyloids in metagenomic samples. A. Normalized abundance of amyloid proteins in gut microbiomes of Parkinson’s disease (PD), Alzheimer’s disease (AD), and cerebral meningitis (CM) patients and controls. The p-values for significant differences (PD patients) are given at the top; where ns stands for non-significant (AD patients). B. Normalized abundance of identified amyloid homologs in PD, AD and CM patients. Red dots represent proteins significantly more abundant in diseased patients compared to the controls (Mann-Whitney U-test with Benjamini-Hochberg correction, p-value < 0.05).

For each protein of the *BFA* database and its homologs from *UGPAmyloids*, we analyzed their individual abundances in each sample to identify protein families contributing most to the observed amyloid enrichment in PD samples (Fig. 3B). A significant correlation between the abundance of biofilm-related CsgA and neurodegeneration could be observed. In two out of three PD studies, homologs of CsgA were found to be more abundant in PD patients (Fig. 3B). Interestingly, the same trend was found in the AD data (Mann-Whitney U-test with Benjamini-Hochberg correction, p-value < 0.05), but not for CM. CsgA is known to interact with PD-related alpha-synuclein and AD-related Abeta peptide, Perov et al. (2019), Bhoite (2022), Otzen et al. (2025). This protein was also found to promote amyloid-related neurodegeneration in *C. elegans* models of AD and PD, Huntington’s disease and Amyotrophic Lateral Sclerosis, Wang et al. (2021). Not all bacterial functional amyloids appear to be equally important from a clinical perspective, and CsgA holds a special place in this regard.

### Identified intestine bacterial functional amyloids are often extracellular and interact with human intestinal proteins

The observed increased abundance of *UHGPAmyloids* in the PD microbiome raises a question about their potential routes of interaction with human cells. The putative pro-inflammatory or toxic effect of bacterial functional amyloids on the human body is closely related to their cellular localization. Bacterial proteins located inside cells are much less likely to come in contact with human proteins and undesired protein-protein interactions.

In general, prokaryotic proteins tend to localize mostly in the cytoplasm (up to 64%), membrane (up to 20%), and only <2% are secreted, Peng and Gao (2014). The analyzed bacterial functional amyloids display, however, different proportions. Firstly, we predicted the cellular localization of the source proteins from the *BFA* dataset, 60% of which belonged to the *Extracellular* and 40% to the *Cytoplasm* class. The *UHGPAmyloids* dataset followed this trend, although the proportions of the predicted cellular compartments were different: *Extracellular* 43.1%, *Cytoplasm* 48.2%, *Plasma membrane* 8.7%. The predicted *Extracellular* proteins are homologs of 25 different amyloids, although the dominant group includes biofilm-related proteins, such as WapA (187 homologs) and Bap (129 homologs).

Protein-protein interactions are central to understanding the molecular mechanisms of stimulus response. The predicted cellular localization provides a basis for analyzing which human proteins might interact with bacterial functional amyloids. From *The Human Protein Atlas*, Uhlén et al. (2015), we extracted proteins that are experimentally confirmed to be expressed in the human intestine and are biologically related to the extracellular membrane or the junction space (*HPAIntestine_filtered*, 2,361 proteins). Similarly, we selected all *UHGPAmyloids* with *Extracellular* or *Plasma Membrane* localization (*UHGPAmyloids_filtered,* 417 proteins). Subsequently, all possible interactions between proteins from *HPAIntestine_filtered* and *UHGPAmyloids_filtered* were computationally predicted.

We found 183,742 interactions involving 1,098 human proteins from *HPAIntestine_filtered.* The distribution of the number of interactors per human protein is skewed (Fig. S2A). For each protein from *UHGPAmyloids_filtered*, there were human interactors predicted. The highest number of human interactors was found for homologs of WapA (114,058 in total) and the lowest for homologs of TapA (135 in total) (Fig. S3B).

The identified human protein potentially interacting with bacterial functional amyloids most abundantly was N-myc-interactor (Nmi protein, UniprotID: Q13287). The analysis showed 681 interactions with homologs from 20 different protein groups, mostly with biofilm-related homologs of WapA (346), SasG (92) and CsgA (70). All of these homologs originate from Firmicutes or Proteobacteria, the latter often seen as a disease marker, Shin et al. (2015). Nmi protein is a part of the human immune system and interacts with interleukin-2 and STAT proteins, Zhu et al. (1999). Its level can be elevated in multiple cancers, He et al. (2022). Among proteins representing the 5 highest numbers of interactions, we also found other immune response-related proteins (C-C motif chemokine 5, Interferon-induced transmembrane protein 1 and 2), claudins - controlling the epithelial barrier and ubiquitin ligases (Supplementary Table 2), Lynn et al. (2020). Interestingly, in this group, we also identified proteins that become human pathological amyloids, mainly alpha-synuclein (417 different interactions), major prion protein (417 interactions), and APP cutting protein - presenilin-1 (417 interactions). Other human amyloids interacting with *UHGPAmyloids* included APP (294 interactions) and tau (229 interactions).

### Bacterial functional amyloids in human intestine affect signaling and transport

To better understand general trends underlying interactions between human intestinal proteins and bacterial functional amyloids, a two-step overrepresentation analysis (ORA) was performed, Wu et al (2021). We extracted proteins from *HPAIntestine_filtered* that are predicted to potentially interact with at least five putative amyloids from *UHGPAmyloids_filtered* (947 proteins). Then, we checked which GO terms are overrepresented with respect to the whole human proteome (Fig. 4, Supplementary Table 3. contains the list of all identified GO/KEGG terms). Finally, we examined if such terms are also overrepresented with respect to the intestinal proteins - *HPAIntestine_filtered* (Supplementary Table 3).

**Fig 4.**
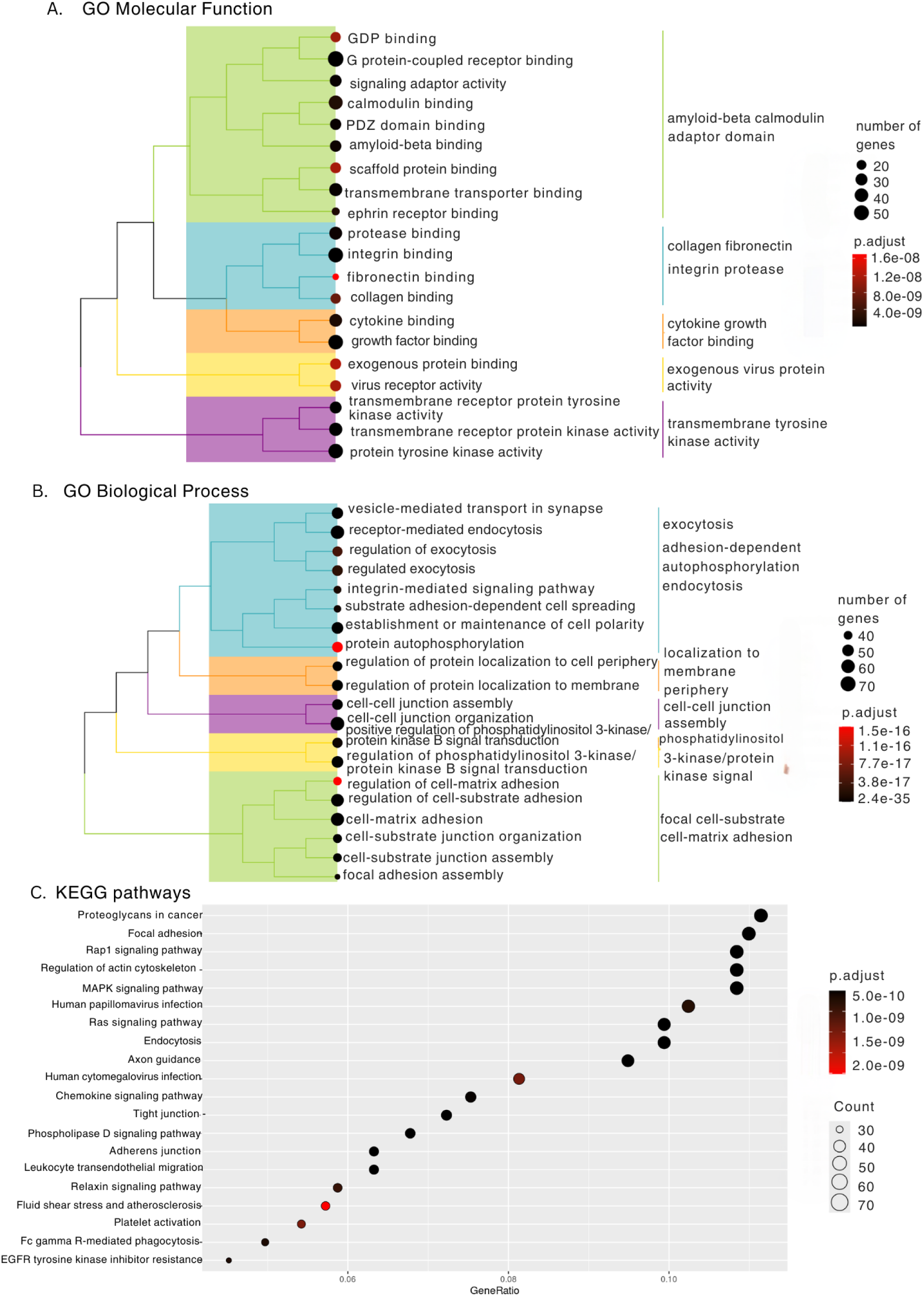
The results of the overrepresentation analysis for A. Gene Ontology Molecular Function, B. Gene Ontology Biological Process C. KEGG pathways.

The overrepresentation analysis revealed that bacterial functional amyloids are not neutral to proteins of the human gastrointestinal tract (Fig. 4). We observed enrichment in GO *Biological Process* terms related to vesicle transport, protein localization, signaling, cell-cell and cell-matrix adhesion. The results showed that identified amyloids are capable of interacting with proteins forming cell-cell junctions, like claudins, which are the main components of Tight Junctions (TJ). This group of proteins is essential for maintaining the integrity of epithelium and is associated with numerous gastrointestinal diseases including Inflammatory Bowel Disease (IBD), Ulcerative Colitis (UC), Crohn’s Disease (CD) or Colorectal Cancer (CRC), Landy et al 2016, Tsukita et al. 2019. Other fundamental human processes, such as signaling and transport, also seem to be affected by bacterial functional amyloids of the gut microbiome. We observed the enrichment in cytokine-binding proteins, growth factors and G-protein-coupled receptors, as well as tyrosine kinase receptors, which all mediate cell response to internal and external factors. Similarly, terms related to endocytosis, exocytosis, and integrin-binding, all regulating protein transport, are also overrepresented. Notably, human pathological amyloids, such as Abeta-42 and tau, hallmarks of Alzheimer’s disease, also bind to integrin and integrin binding receptors, Wang and Ye (2020), Venkatasubramaniam et al. (2014), which suggests that some interactors are shared by bacterial functional and human pathological amyloids.

Analogously, overrepresentation results concerning KEGG pathways (Fig. 4C) and Disease Ontology (Fig. S3) were analyzed. Apart from pathways related to cell-to-cell adhesion, including focal adhesion, cell and adherens junctions, or transport, multiple cancer-related pathways seem to be affected. The most profound effect can be observed for the pathway “Proteoglycans in cancer”. This pathway concerns proteins in the microenvironment of the cancerous cells affecting numerous processes, such as cell proliferation, migration, and adhesion. It is also linked to the MAPK signaling pathway, which plays an important role in cell proliferation, differentiation and cell death. In addition, the disease ontology results (Supplementary, Fig. S4) also indicate a significant population of proteins related to cancer disorders. Interestingly, bacterial functional amyloids can potentially bind to human proteins involved in viral infections, as well as other immunity-related pathways such as chemokine signaling pathway and leukocyte migration.

Finally, we evaluated if the predicted human interactors of the bacterial functional amyloids are also the interactors of human amyloid proteins. For this goal, we extracted the Amyloid Interactome published in Biza et al. (2017). They screened protein-protein interaction databases to identify a dataset of human proteins that interact with human amyloids, such as pathological alpha-synuclein, prion protein or tau. We predicted that 61 interactors of the human amyloid proteins, out of 251 present in the Amyloid Interactome, interact with bacterial functional amyloids. Fisher’s exact test for overrepresentation with p-value<10^-6^ suggested a statistically significant overrepresentation of Amyloid Interactome proteins among human proteins predicted to interact with bacterial functional amyloids.

## Discussion

Growing evidence on the impact of the human gut microbiome on the onset and progression of neurological diseases naturally raised questions about mechanisms governing this relation. Bidirectional interplay between the gut microbiota and the central nervous system can occur via the enteric nervous system or immunological response mechanisms, interactions between neuroactive molecules derived from enteric microbiota with neuronal, immune or neuroendocrine gut-derived proteins, Forsyth et al. (2011), Heravi et al. (2023), Kelly et al. (2013), Mirzaei et al. (2021), Mayer et al. (2022).

Recent studies suggested a potential role for certain bacterial functional amyloids in this arena, Hossain et al. (2019), Jeong et al. (2019), Shin et al. (2015). These proteins can be expressed by bacteria inhabiting human intestines and show intriguing structural similarity to misfolded human pathological amyloids that mark neurodegeneration. Additionally, as it turns out, little homology is not an obstacle to form stable structural contacts between such human and bacterial proteins, Otzen at al. (2025). Based on the studies on PD, a few potential modes of action of the bacterial functional amyloids and their contribution to neurodegeneration could be proposed. Firstly, in the light of the Braak hypothesis that PD is initiated in the enteric nervous system and then propagates to the brain via the vagus nerve, Braak et al. (2003), bacterial functional amyloids could serve as triggers of the pathological aggregation of human alpha-synuclein in colon neurons via direct protein-protein interactions, such as cross-seeding, Elkins et al. (2024). Secondly, the seeds from bacterial amyloids were recently shown to be able to propagate across the endothelial barriers through opening cadherin-based junctions (Shenoy, 2019) and enter brain through systemic circulation, despite the blood-brain barrier (Otzen et al. 2025). Notably, these junctions open in the transport of CsgA protein across the gastric wall (Otzen et al. 2025 and references therein), which may indicate a significant role of this protein in PD. Finally, their remarkable structural similarity to human pathological amyloids could activate similar faulty molecular pathways, in particular including immune-related ones Bucianntini et al. (2002). For example, both pathological and functional amyloids are capable of activating Toll-like receptors and signal inhibitory receptors on leukocytes-1 (SIRL-1), hence modulating the immune response Golan et al. (2022).

To investigate these hypotheses, we provided the first analysis of the landscape of bacterial functional amyloids produced by the human gut microbiome, and assessed their potential impact on the onset of neurodegenerative diseases.

We identified a significantly diverse set of putative bacterial functional amyloids in the human gut proteome via homology search in seven bacterial phyla, highlighting that the known bacterial functional amyloids may be just a tip of an iceberg.

To support the conjecture concerning the potential role of these proteins in the disease, based on the publicly available metagenomic datasets, we showed that microbiomes of PD patients across all publicly available studies contain more bacterial functional amyloids than their healthy controls. Notably, the total abundance of bacterial functional amyloids in AD and CM patients did not show significant differences from the respective control groups (Fig. 3). Although the overall greater abundance of gut bacterial functional amyloids was not shown to be universal for both neurodegenerative diseases, the abundance of the specific protein family, CsgA, was. We also observed a greater abundance of CsgA homologs in positive AD, CM and 2 out of 3 PD cohorts. The significant differences of this protein abundances further confirm its potentially salient role in development and progression of neurodegenerative diseases, utilizing the discovered modes of action (Otzen et. al, 2025). It is conceivable that the structural similarity between functional bacterial amyloids and pathological human amyloids could enable molecular mimicry, leading to enhanced aggregation of pathological amyloids or the activation of the same molecular pathways, Golan et al. (2022), Burdukiewicz et al. (2023). It was discovered that CsgA could interact with alpha-synuclein, as well as with Abeta, *in vivo* and *in vitro*, and inhibition of CsgA aggregation reduces neuronal death in *C. elegans* model study due to cross-seeding, Perov et al. (2019), Wang et al. (2021), Bhoite (2022). Moreover, it has been reported that CsgA amyloid precursor contains a pathogen-associated molecular pattern (PAMP) that is recognized by the human immune system in the same manner as Abeta-42 peptide, due to their structural features and despite different sequences of amino acids. CsgA, similarly as Abeta-42, are recognized by TLR2/TLR1 immune sensor-receptor system consisting of 13 different TLR-type receptors that control the same up-regulation of IL-17A- and IL-22-mediated pro-inflammatory-signaling pathways, Zhou et al. (2012), Hill and Lukiw (2015). Hence, the observed greater abundance of CsgA alone may facilitate the onset of AD by activating inflammation and activity of cytokines, deteriorating vascular permeability and levels of free-radicals, as well as enhancing the fibrillation of human amyloidogenic proteins. All of these factors are known to contribute to neurodegeneration.

Accordingly, the interactions between bacterial functional amyloids and human proteins could have a much broader scale. The amyloid proteins found in our study (*UHGPAmyloids*), are 20-fold more likely to be extracellular than other bacterial proteins, according to the cellular localization predictions, which potentially enable them for molecular interactions with human proteins. Following our analyses, such direct interactions may affect proteins responsible for endo-and exocytosis, signaling and cellular transport.

The bacterial functional amyloids could be one of the factors contributing to pro-inflammatory characteristics of certain phyla. For example, Proteobacteria that produce CsgAB turned out to be rich in such proteins. The increased population of Proteobacteria was observed in Parkinson’s disease, type II diabetes, and Alzheimer’s disease, Ali Keshavarzian et al. (2015), Zhang et al. (2021), Hung et al. (2022), Nagpal et al. (2019). On the other hand, we also found amyloids in anti-inflammatory bacteria, like *Lactobacillus* and *Bifidobacterium*. Although the number of amyloid proteins in these bacteria was low, it could be an important observation due to their frequent clinical applications, which may be relevant in cases of defective gut permeability.

In addition, the bacterial functional amyloids, similar to human pathogenic amyloids, seem to have the potential to affect cell junctions, Yamazaki et al. (2019). The negative effect of the protein aggregation phenomena on epithelial cell integrity has been already observed, Yamazaki et al. (2019). The amount of tight junctions, necessary e.g. for proper blood-brain barrier functionality, has been shown to negatively correlate with Abeta, tau and apolipoprotein abundances, Yamazaki et al. (2019), Liu et al. (2020). Moreover, alpha-synuclein and Abeta were found to disrupt the expression of tight junctions and their downregulation, Yamazaki et al., (2019), Bruban et al. (2009), Kuan et al. (2016), Marco and Scaper (2006). Therefore, based on these observations, the abundance of bacterial functional amyloids in the gut may influence the gut permeability, similarly as human pathological amyloids affect the blood-brain barrier.

According to our results, bacterial functional amyloids have the potential to induce inflammatory responses. They are capable of interacting with immune response proteins involved in chemokine signaling and leukocyte migration. Bacterial functional amyloids produced by gut bacteria, similar to human pathological amyloids, could bind integrin and calmodulin. Abeta and tau are capable of binding to integrin, which leads to neurotoxic effect, Wright et al. (2007), Wang & Ye (2021). Calmodulin binding to neuroinflammatory proteins, like alpha-synuclein, Abeta and apolipoprotein, correlates with neurodegenerative events, O’Day and Huber (2022).

Our results demonstrate that bacterial functional amyloids are other important actors in the gut-brain axis. The specific microbiome composition observed in PD has altered functionality caused, e.g., by the increase of bacterial species expressing functional amyloids. This can lead to a pro-inflammatory positive feedback loop, in which bacterial functional amyloids affect immune response and interact with tight junctions, inciting a greater gut permeability. This, in turn, enables them to penetrate the epithelial barrier and promote further inflammation. Finally, the potential of bacterial functional amyloids to interact with similar proteins as human pathological amyloids, along with their structural resemblance, may induce alpha-synuclein aggregation in the enteric nervous system. This aggregation could then propagate via the vagus nerve to the brain, aligning with the Braak hypothesis, Braak et al. (2003). The presented framework of molecular amyloid interactions in PD, although focused on one set of gut-derived molecules, could open the door for novel research strategies in neurodegeneration but requires further *in vivo* experiments.

## Data availability

The datasets produced in this paper (including *UHGPAmyloids* and predicted protein-protein interactions) are available on a Zenodo repository: https://zenodo.org/records/14016809

## Funding

This work was supported by the National Science Centre, Poland [2019/35/B/NZ2/03997] (MK) and NAWA STER grant of Polish National Agency for Academic Exchange for PhD students (JWW). TK is funded by the National Science Centre, Poland [2023/05/Y/NZ2/00080].

## Acknowledgements

We would like to cordially thank Prof. Pawel P. Łabaj for valuable comments and a fruitful discussion on the manuscript.

Access to Wroclaw Centre for Networking and Supercomputing is greatly acknowledged.

## Methods

### Identification of amyloids in human gut

To identify putative amyloids in the human gut, we used two datasets: Unified Human Gut Proteome *UHGP* from Almeida et al. (2020) and a set of bacterial functional amyloids *BFA* produced in Nowakowska et al. (2023). Unified Human Gut Proteome project provides multiple datasets. We used a version of *UHGP* clustered on 95% of sequence identity in a v1.0 edition (file name: uhgp-95.faa). The dataset was downloaded from the website http://ftp.ebi.ac.uk/pub/databases/metagenomics/mgnify_genomes/human-gut/v1.0/.

We searched for homologs of *BFA* in *UHGP* using mmseqs with the command: mmseqs search bfa uhgp-95 results tmp --comp-bias-corr 0 --mask 0 Mirdita et al. (2019). The lack of composition correction was purposeful. Multiple functional amyloids contain tandem repeats and low complexity regions which are responsible for the aggregation, Nowakowska et al. (2023). Hence, we did not mask such regions during homology search.

We filtered the amyloids using AMYPred-FRL software, available as a web server, with the cutoff value of 0.8 Charoenkwan et al. (2022). AMYPred-FRL is an entire sequence-based predictor. To further examine the aggregation potential of our sequences we used ArchCandy, Ahmed et al. (2015). For each sequence, we used a cutoff value 0.5 and ran the following command: java-jar ArchCandyV2_CLI.jar –TMfilter-t=0.5-i=Seq_ID Sequence.

The dataset of homologs of *BFA* in *UHGP* with positive aggregation predictions was further denoted as *UHGPAmyloids*.

Clusterization of *UHGPAmyloids* sequences was performed with CD-HIT using the following command: cd-hit-i input_file.fasta-o output_file.fasta-c 0.9/0.8/0.7/. Fu et al. (2012).

### Taxonomic analysis

Taxonomic assignments of *UHGPAmyloids* sequences were already provided in the *UHGP* dataset under the file genomes-all_metadata.tsv. The taxonomic nomenclature throughout the text is consistent with the one provided in this file. Shannon entropy calculations were performed with the R *phyloseq* package designed for taxonomic analyses, McMurdi and Holmes (2013).

### Metagenomic analysis

To investigate the occurrence of functional amyloids in amyloid-related disease patients, there were analyzed data from three Parkinson’s disease studies, Bedarf et al. (2017), Wallen et al. (2022), Boktor et al. (2023), one Alzheimer’s disease study, Laske et al. (2022), and Cryptococcal Meningitis study, Li et al. 2023. Raw reads for the metagenomics section of the analysis were downloaded from the Sequence Read Archive (SRA). All samples were quality-controlled using Trim Galore 0.6.10 (Kruger 2015) with default parameters. Quality-controlled reads were then assembled using MetaSpades 3.15.5 with default parameters, Nurk et al. 2017. Genes were predicted using Prodigal 2.6.3, Hyatt et al. 2010.

Then, to estimate coverage, reads were mapped into predicted genes using KMA 1.4.14, Clausen et al. 2018. To estimate relative abundance, the number of reads mapped to each gene was divided by the sum of all mapped reads. Previously identified amyloids from *UHGPAmyloids* were then searched in all samples using MMseqs2 15.6f452 with default parameters, Steinegger and Soeding 2017. Obtained hits were then filtered by percentage of identical residues (>90) to ensure that only highly similar sequences were analyzed (see Fig. 5).

**Fig. 5.**
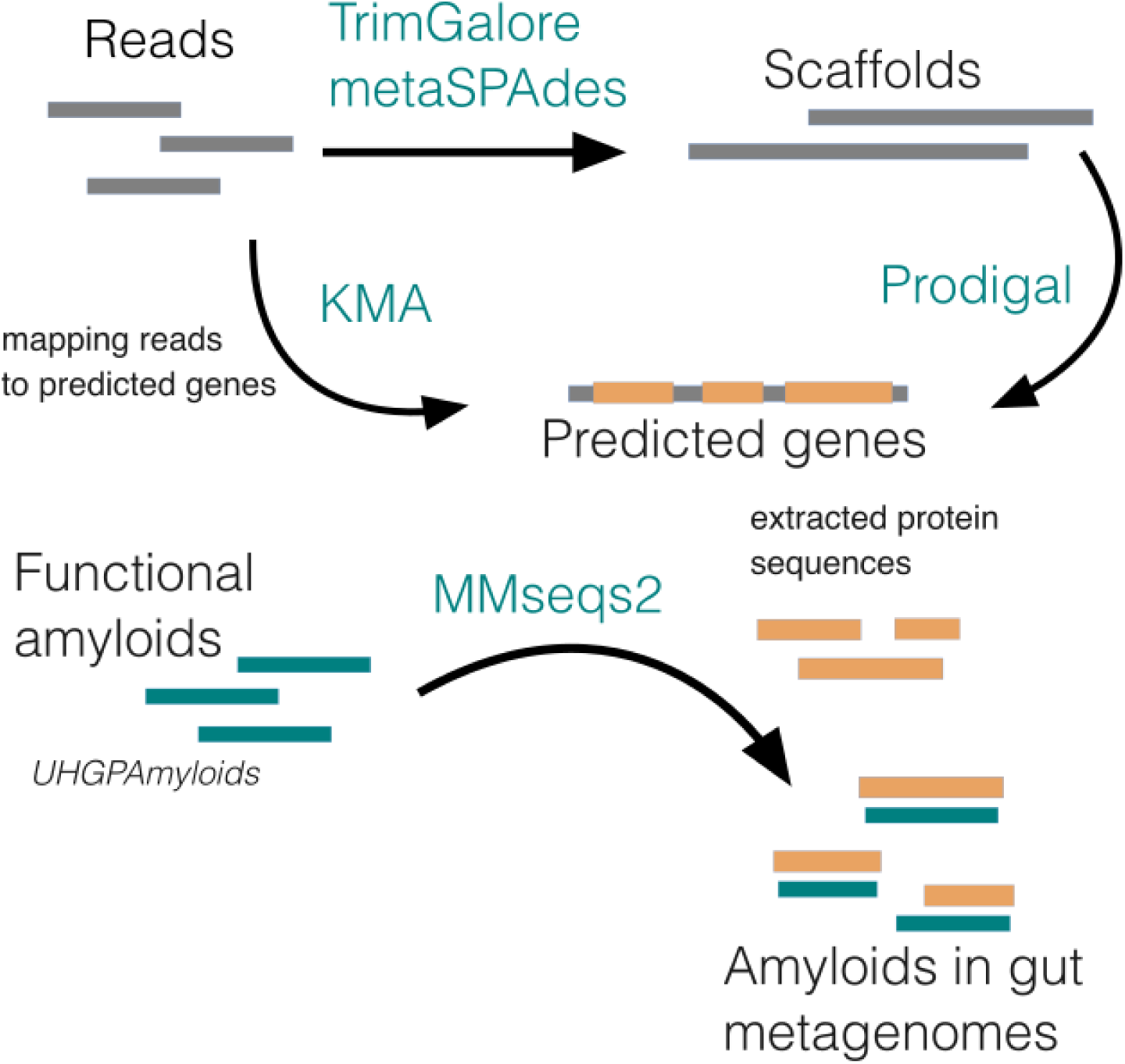
The pipeline used for identification of amyloids in metagenomic samples.

To test the difference in abundance of amyloids between case and control groups, Mann-Whitney U-test was applied. For comparing abundances of specific proteins between case and control groups, Mann-Whitney U-test with Benjamini-Hochberg correction was applied, Benjamini and Hochberg (1995).

### Prediction of cellular localization

To predict cellular localization of a protein, we used BUSCA, Savojardo et al. (2018). BUSCA predictions are based on identification of different peptides and domains. All predicted amyloids were run on the web server with a taxonomic group: Prokarya - Other - 3 compartments.

### Protein-protein interaction prediction

For protein-protein interaction prediction, we only considered proteins from *UHGPAmyloids* and *Extracellular* or *Plasma membrane* localization according to BUSCA. The human proteins were extracted from *The Human Protein Atlas* Uhlén et al. (2015), file normal_tissue.tsv (version: 23.0) available in the Downloadable data section on the website. normal_tissue.tsv was filtered to contain only proteins expressed in the intestine. To do so, we considered proteins that contained the colon or small intestine or duodenum or rectum in the *Tissue* column. Next, we filtered the dataset with respect to the cellular localization. For each intestine protein, we extracted its Subcellular location [CC] and checked if it contained an expression from the following set: {Cell membrane, cell membrane, secreted, Secreted, extracellular, Extracellular, cell surface, Cell surface, junction, Junction, secretory, Secretory, cell wall, Cell wall}. This way we identified 2361 proteins that are potentially at the first line of contact with substances secreted by the microbiome, which were further denoted as *HPAIntestine_filtered*.

We performed predictions of protein-protein interactions with ProteinPrompt, Canzler et al. (2022) (using default parameters) between proteins from *UHGPAmyloids with Extracellular or Membrane* cellular localization according to BUSCA *(UHGPAmyloids_filtered)* and *HPAIntestine_filtered* which can be defined as:

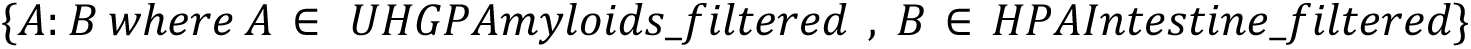

The overrepresentation analysis was performed for those proteins from *HPAIntestine_filtered* that had at least 5 interacting partners in *UHGPAmyloids_filtered.* The identification of enriched terms was done with the ClusterProfiler package available in R, Wu et al. (2021) (version: 4.10). The specific functions used there were: *enrichGO*, *enrichKEGG* and *enrichDO*.

The F-test was performed for top 20 terms for *Biological Process*, *Molecular Function* and *KEGG* between proteins *HPAIntestine_filtered* which had at least 5 interacting partners in *UHGPAmyloids* and the entire set *HPAIntestine_filtered* with *f.test* function in R programming language. Each p-value was multiplied by 60, in accordance with the Bonferroni correction.

### General data analysis

All downstream analyses were performed using the Python programming language with NumPy 1.24.3, Harris et al. 2020, Pandas 2.0.3, McKinney 2011, SciPy 1.11.1, Virtanen et al. 2020, and Matplotlib 3.7.2, Hunter 2007 packages. Additional visualizations were performed in R using ggplot2 3.5.1, Wickham (2014).

## Notes

https://zenodo.org/records/14016809

### Competing Interest Statement

The authors have declared no competing interest.

### Summary of Updates

added paragraphs to the discussion and the results

## References

Ahmed, Abdullah B., et al. “A structure-based approach to predict predisposition to amyloidosis.” Alzheimer’s & Dementia 11.6 (2015): 681–690.

Almeida, Alexandre, et al. “A unified catalog of 204,938 reference genomes from the human gut microbiome.” Nature biotechnology 39.1 (2021): 105–114.

Bedarf, Janis R., et al. “Functional implications of microbial and viral gut metagenome changes in early stage L-DOPA-naïve Parkinson’s disease patients.” Genome medicine 9 (2017): 1–13.

Biza, Konstantina V., et al. “The amyloid interactome: Exploring protein aggregation.” PloS one 12.3 (2017): e0173163.

Bhoite, Sujeet S., et al. “Mechanistic insights into accelerated α-synuclein aggregation mediated by human microbiome-associated functional amyloids.” Journal of Biological Chemistry 298.7 (2022).

Boktor, Joseph C., et al. “Integrated Multi-Cohort Analysis of the Parkinson’s Disease Gut Metagenome.” Movement Disorders 38.3 (2023): 399–409.

Braak, Heiko, et al. “Idiopathic Parkinson’s disease: possible routes by which vulnerable neuronal types may be subject to neuroinvasion by an unknown pathogen.” Journal of neural transmission 110 (2003): 517–536.

Bruban, Julien, et al. “Amyloid-β (1-42) alters structure and function of retinal pigmented epithelial cells.” Aging cell 8.2 (2009): 162–177.

Bucciantini, Monica, et al. “Inherent toxicity of aggregates implies a common mechanism for protein misfolding diseases.” nature 416.6880 (2002): 507–511.

Burdukiewicz, Michał, et al. “Amylograph: a comprehensive database of amyloid–amyloid interactions.” Nucleic Acids Research 51.D1 (2023): D352–D357.

Cattaneo, Annamaria, et al. “Association of brain amyloidosis with pro-inflammatory gut bacterial taxa and peripheral inflammation markers in cognitively impaired elderly.” Neurobiology of aging 49 (2017): 60–68.

Canzler, Sebastian, et al. “ProteinPrompt: a webserver for predicting protein–protein interactions.” Bioinformatics advances 2.1 (2022): vbac059.

Charoenkwan, Phasit, et al. “AMYPred-FRL is a novel approach for accurate prediction of amyloid proteins by using feature representation learning.” Scientific reports 12.1 (2022): 7697.

Chen, Lili, et al. “A comparison of the composition and functions of the oral and gut microbiotas in Alzheimer’s patients.” Frontiers in Cellular and Infection Microbiology 12 (2022): 942460.

Clausen, Philip TLC, Frank M. Aarestrup, and Ole Lund. “Rapid and precise alignment of raw reads against redundant databases with KMA.” BMC bioinformatics 19 (2018): 1–8.

Dodiya, Hemraj B., et al. “Synergistic depletion of gut microbial consortia, but not individual antibiotics, reduces amyloidosis in APPPS1-21 Alzheimer’s transgenic mice.” Scientific reports 10.1 (2020): 8183.

Dueholm, Morten S., Daniel Otzen, and Per Halkjær Nielsen. “Evolutionary insight into the functional amyloids of the pseudomonads.” PLoS One 8.10 (2013): e76630.

Dueholm, Morten S., et al. “Curli functional amyloid systems are phylogenetically widespread and display large diversity in operon and protein structure.” PloS one 7.12 (2012): e51274.

Elkins, Molly, Neha Jain, and Çagla Tükel. “The menace within: bacterial amyloids as a trigger for autoimmune and neurodegenerative diseases.” Current Opinion in Microbiology 79 (2024): 102473.

Fang, P., et al. “The microbiome as a modifier of neurodegenerative disease risk.” Cell host & microbe 28.2 (2020): 201–222.

Friedland, Robert P., and Matthew R. Chapman. “The role of microbial amyloid in neurodegeneration.” PLoS pathogens 13.12 (2017): e1006654.

Friedland, Robert P. “Mechanisms of molecular mimicry involving the microbiota in neurodegeneration.” Journal of Alzheimer’s Disease 45.2 (2015): 349–362.

Fu, Limin, et al. “CD-HIT: accelerated for clustering the next-generation sequencing data.” Bioinformatics 28.23 (2012): 3150–3152.

Fusco, William, et al. “Short-chain fatty-acid-producing bacteria: key components of the human gut microbiota.” Nutrients 15.9 (2023): 2211.

Gallardo, Rodrigo, Neil A. Ranson, and Sheena E. Radford. “Amyloid structures: much more than just a cross-β fold.” Current opinion in structural biology 60 (2020): 7–16.

Gilbert, Jack A., et al. “Current understanding of the human microbiome.” Nature medicine 24.4 (2018): 392–400.

Golan, Nimrod, Yizhaq Engelberg, and Meytal Landau. “Structural mimicry in microbial and antimicrobial amyloids.” Annual Review of Biochemistry 91.1 (2022): 403–422.

Harach, Taoufiq, et al. “Reduction of Abeta amyloid pathology in APPPS1 transgenic mice in the absence of gut microbiota.” Scientific reports 7.1 (2017): 41802.

Harris, Charles R., et al. “Array programming with NumPy.” Nature 585.7825 (2020): 0352,357-362.

He, Teng, et al. “NMI: a potential biomarker for tumor prognosis and immunotherapy.” Frontiers in pharmacology 13 (2022): 1047463.

Heravi, Fatemah Sadeghpour, Kaveh Naseri, and Honghua Hu. “Gut microbiota composition in patients with neurodegenerative disorders (Parkinson’s and Alzheimer’s) and healthy controls: a systematic review.” Nutrients 15.20 (2023): 4365.

Hill, James M., and Walter J. Lukiw. “Microbial-generated amyloids and Alzheimer’s disease (AD).” Frontiers in aging neuroscience 7 (2015): 9.

Hirayama, Masaaki, and Kinji Ohno. “Parkinson’s disease and gut microbiota.” Annals of Nutrition and Metabolism 77.Suppl. 2 (2021): 28–35.

Hung, Chun-Che, et al. “Gut microbiota in patients with Alzheimer’s disease spectrum: A systematic review and meta-analysis.” Aging (Albany NY) 14.1 (2022): 477.

Hunter, John D. “Matplotlib: A 2D graphics environment.” Computing in science & engineering 9.03 (2007): 90–95.

Hyatt, Doug, et al. “Prodigal: prokaryotic gene recognition and translation initiation site identification.” BMC bioinformatics 11 (2010): 1–11.

Ivanov, Ivaylo I., et al. “Induction of intestinal Th17 cells by segmented filamentous bacteria.” Cell 139.3 (2009): 485–498.

Kelly, Leo P., et al. “Progression of intestinal permeability changes and alpha-synuclein expression in a mouse model of Parkinson’s disease.” Movement Disorders 29.8 (2014): 999–1009.

Keshavarzian, Ali, et al. “Colonic bacterial composition in Parkinson’s disease.” Movement Disorders 30.10 (2015): 1351–1360.

Knight, Rob, et al. “The microbiome and human biology.” Annual review of genomics and human genetics 18.1 (2017): 65–86.

Konstantoulea, Katerina, et al. “Heterotypic interactions in amyloid function and disease.” The FEBS Journal 289.8 (2022): 2025–2046.

Kuan, Wei-Li, et al. “α-Synuclein pre-formed fibrils impair tight junction protein expression without affecting cerebral endothelial cell function.” Experimental neurology 285 (2016): 72–81.

Landy, Jonathan, et al. “Tight junctions in inflammatory bowel diseases and inflammatory bowel disease associated colorectal cancer.” World journal of gastroenterology 22.11 (2016): 3117.

Laske, Christoph, et al. “Signature of Alzheimer’s disease in intestinal microbiome: results from the AlzBiom study.” Frontiers in Neuroscience 16 (2022): 792996.

LeBlanc, Jean Guy, et al. “Bacteria as vitamin suppliers to their host: a gut microbiota perspective.” Current opinion in biotechnology 24.2 (2013): 160–168.

Lebouvier, Thibaud, et al. “Colonic biopsies to assess the neuropathology of Parkinson’s disease and its relationship with symptoms.” PloS one 5.9 (2010): e12728.

Levkovich, Shon A., Ehud Gazit, and Dana Laor Bar-Yosef. “Two decades of studying functional amyloids in microorganisms.” Trends in Microbiology 29.3 (2021): 251–265.

Liu, Chia-Chen, et al. “Tau and apolipoprotein E modulate cerebrovascular tight junction integrity independent of cerebral amyloid angiopathy in Alzheimer’s disease.” Alzheimer’s & Dementia 16.10 (2020): 1372–1383.

Lynn, K. Sabrina, Raven J. Peterson, and Michael Koval. “Ruffles and spikes: Control of tight junction morphology and permeability by claudins.” Biochimica et Biophysica Acta (BBA)-Biomembranes 1862.9 (2020): 183339.

Marco, Sonia, and Stephen D. Skaper. “Amyloid β-peptide1–42 alters tight junction protein distribution and expression in brain microvessel endothelial cells.” Neuroscience letters 401.3 (2006): 219–224.

Mayer, Emeran A., Karina Nance, and Shelley Chen. “The gut–brain axis.” Annual review of medicine 73.1 (2022): 439–453.

McKinney, Wes. “pandas: a foundational Python library for data analysis and statistics.” Python for high performance and scientific computing 14.9 (2011): 1–9.

McMurdie, Paul J., and Susan Holmes. “phyloseq: an R package for reproducible interactive analysis and graphics of microbiome census data.” PloS one 8.4 (2013): e61217.

Mirdita, Milot, Martin Steinegger, and Johannes Söding. “MMseqs2 desktop and local web server app for fast, interactive sequence searches.” Bioinformatics 35.16 (2019): 2856–2858.

Nowakowska, Alicja W., et al. “The role of tandem repeats in bacterial functional amyloids.” Journal of Structural Biology 215.3 (2023): 108002.

Nurk, Sergey, et al. “metaSPAdes: a new versatile metagenomic assembler.” Genome research 27.5 (2017): 824–834.

Otoo, Henry N., et al. “Candida albicans Als adhesins have conserved amyloid-forming sequences.” Eukaryotic cell 7.5 (2008): 776–782.

Otzen, Daniel, and Roland Riek. “Functional amyloids.” Cold Spring Harbor perspectives in biology 11.12 (2019): a033860.

Otzen, Daniel E., et al. “Interactions between pathological and functional amyloid: A match made in Heaven or Hell?.” Molecular Aspects of Medicine 103 (2025): 101351.

O’Day, Danton H., and Robert J. Huber. “Calmodulin binding proteins and neuroinflammation in multiple neurodegenerative diseases.” BMC neuroscience 23.1 (2022): 10.

Palanikumar, Loganathan, et al. “Protein mimetic amyloid inhibitor potently abrogates cancer-associated mutant p53 aggregation and restores tumor suppressor function.” Nature communications 12.1 (2021): 3962.

Peng, Chong, and Feng Gao. “Protein localization analysis of essential genes in prokaryotes.” Scientific reports 4.1 (2014): 6001.

Sampson, Timothy R., et al. “A gut bacterial amyloid promotes α-synuclein aggregation and motor impairment in mice.” elife 9 (2020): e53111.

Savojardo, Castrense, et al. “BUSCA: an integrative web server to predict subcellular localization of proteins.” Nucleic acids research 46.W1 (2018): W459–W466.

Scheperjans, Filip, et al. “Gut microbiota are related to Parkinson’s disease and clinical phenotype.” Movement Disorders 30.3 (2015): 350–358.

Schirmer, Melanie, et al. “Linking the human gut microbiome to inflammatory cytokine production capacity.” Cell 167.4 (2016): 1125–1136.

Shannon, Kathleen M., et al. “Alpha-synuclein in colonic submucosa in early untreated Parkinson’s disease.” Movement Disorders 27.6 (2012): 709–715.

Shin, Na-Ri, Tae Woong Whon, and Jin-Woo Bae. “Proteobacteria: microbial signature of dysbiosis in gut microbiota.” Trends in biotechnology 33.9 (2015): 496–503.

Steinegger, Martin, and Johannes Söding. “MMseqs2 enables sensitive protein sequence searching for the analysis of massive data sets.” Nature biotechnology 35.11 (2017): 1026–1028.

Szulc, Natalia, et al. “Variability of amyloid propensity in imperfect repeats of csga protein of salmonella enterica and escherichia coli.” International journal of molecular sciences 22.10 (2021): 5127.

Tsukita, Sachiko, Hiroo Tanaka, and Atsushi Tamura. “The claudins: from tight junctions to biological systems.” Trends in biochemical sciences 44.2 (2019): 141–152.

Uhlén, Mathias, et al. “Tissue-based map of the human proteome.” Science 347.6220 (2015): 1260419.

Vander Borght, Thierry, et al. “Cerebral Metabolic Differences in Parkinson’s and Alzheimer’s Diseases Matched for Dementia Severity.” Journal of Nuclear Medicine 38.5 (1997): 797–802.

Venkatasubramaniam, Arundhathi, Alexander Drude, and Theresa Good. “Role of N-terminal residues in Aβ interactions with integrin receptor and cell surface.” Biochimica et Biophysica Acta (BBA)-Biomembranes 1838.10 (2014): 2568-2577.

Virtanen, Pauli, et al. “SciPy 1.0: fundamental algorithms for scientific computing in Python.” Nature methods 17.3 (2020): 261–272.

Wallen, Zachary D., et al. “Metagenomics of Parkinson’s disease implicates the gut microbiome in multiple disease mechanisms.” Nature communications 13.1 (2022): 6958.

Wang, Chenyin, et al. “Genome-wide screen identifies curli amyloid fibril as a bacterial component promoting host neurodegeneration.” Proceedings of the National Academy of Sciences 118.34 (2021): e2106504118.

Wang, Peng, and Yihong Ye. “Filamentous recombinant human Tau activates primary astrocytes via an integrin receptor complex.” Nature Communications 12.1 (2021): 95.

Wang, Su-shan, et al. “The relationship between Alzheimer’s disease and intestinal microflora structure and inflammatory factors.” Frontiers in aging neuroscience 14 (2022): 972982.

Wickham, Hadley, Winston Chang, and Maintainer Hadley Wickham. “Package ‘ggplot2’.” Create elegant data visualisations using the grammar of graphics. Version 2.1 (2016): 1–189.

Wright, Sarah, et al. “α2β1 and αVβ1 integrin signaling pathways mediate amyloid-β-induced neurotoxicity.” Neurobiology of aging 28.2 (2007): 226–237.

Wu, Tianzhi, et al. “clusterProfiler 4.0: A universal enrichment tool for interpreting omics data.” The innovation 2.3 (2021).

Yamazaki, Yu, et al. “Selective loss of cortical endothelial tight junction proteins during Alzheimer’s disease progression.” Brain 142.4 (2019): 1077–1092.

Yu, Guangchuang, et al. “clusterProfiler: an R package for comparing biological themes among gene clusters.” Omics: a journal of integrative biology 16.5 (2012): 284–287.

Zeng, M. Y., N. Inohara, and G. Nuñez. “Mechanisms of inflammation-driven bacterial dysbiosis in the gut.” Mucosal immunology 10.1 (2017): 18–26.

Zhang, Zewen, et al. “Characteristics of the gut microbiome in patients with prediabetes and type 2 diabetes.” PeerJ 9 (2021): e10952.

Zhou, Yizhou, et al. “Bacterial amyloids.” Amyloid Proteins: Methods and Protocols (2012): 303–320.

Zhu, Ming-hua, et al. “Functional association of Nmi with Stat5 and Stat1 in IL-2-and IFN γ-mediated signaling.” Cell 96.1 (1999): 121–130.

