## Supplementary for "Aggregating gut: on the link between neurodegeneration and bacterial functional amyloids"

#### Figures

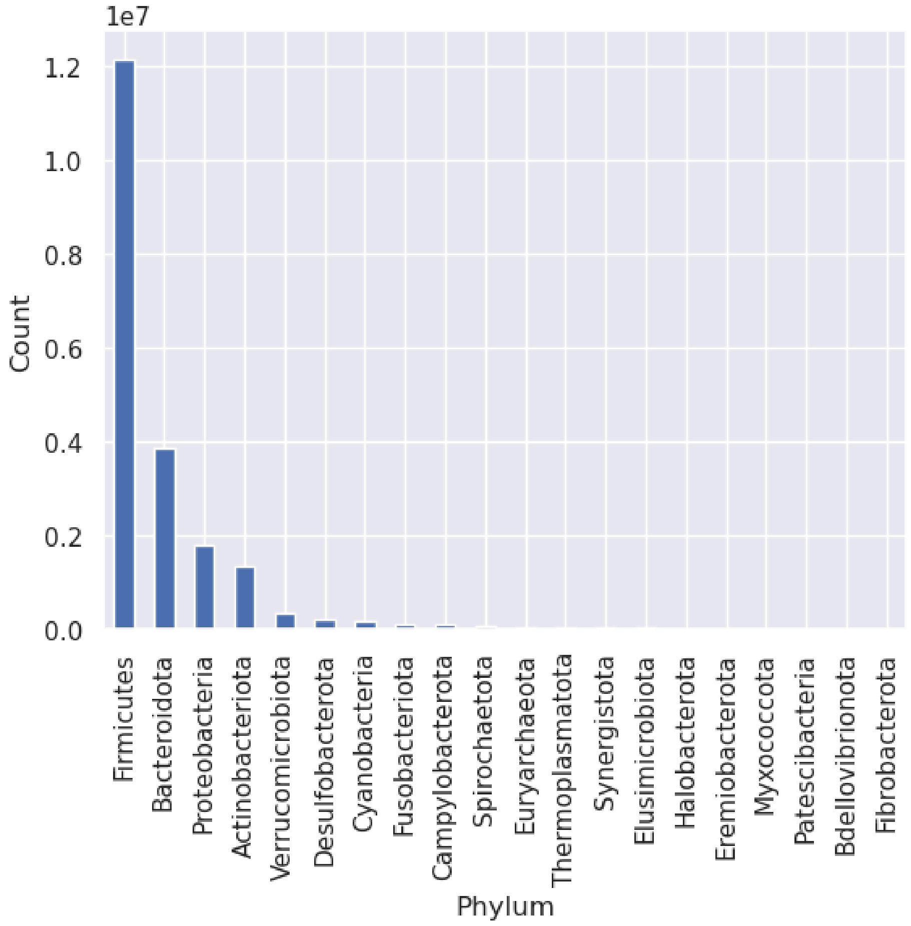


Figure S1. Abundance of different phyla in *UHGP* dataset.

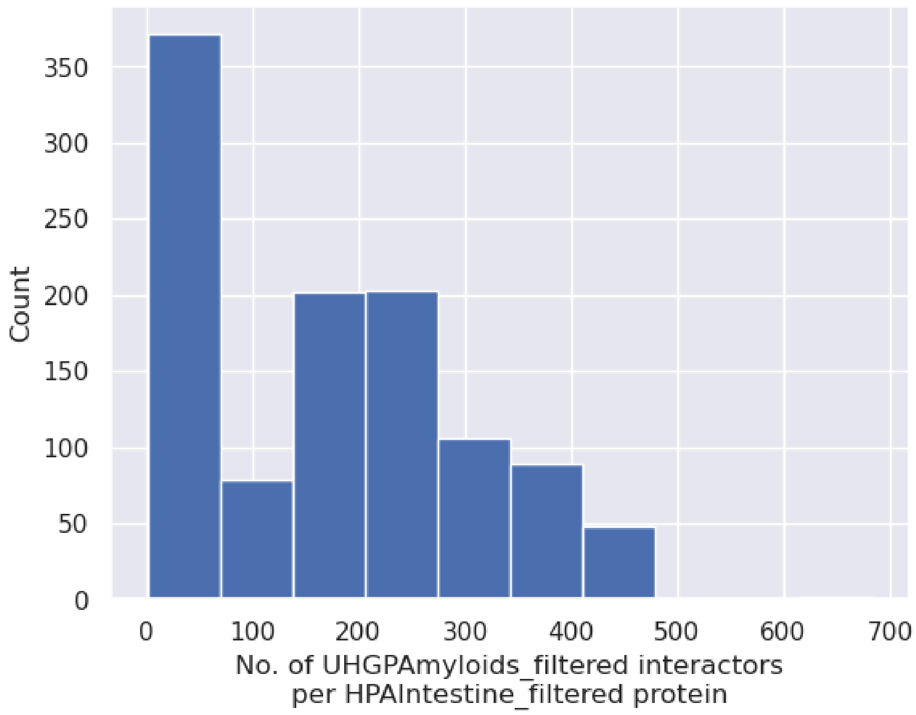


Fig S2. Distribution of the number of interactors from *UHGPAmyloids_filtered* per protein from *HPAintestine_filtered* based on protein-protein interaction predictions.

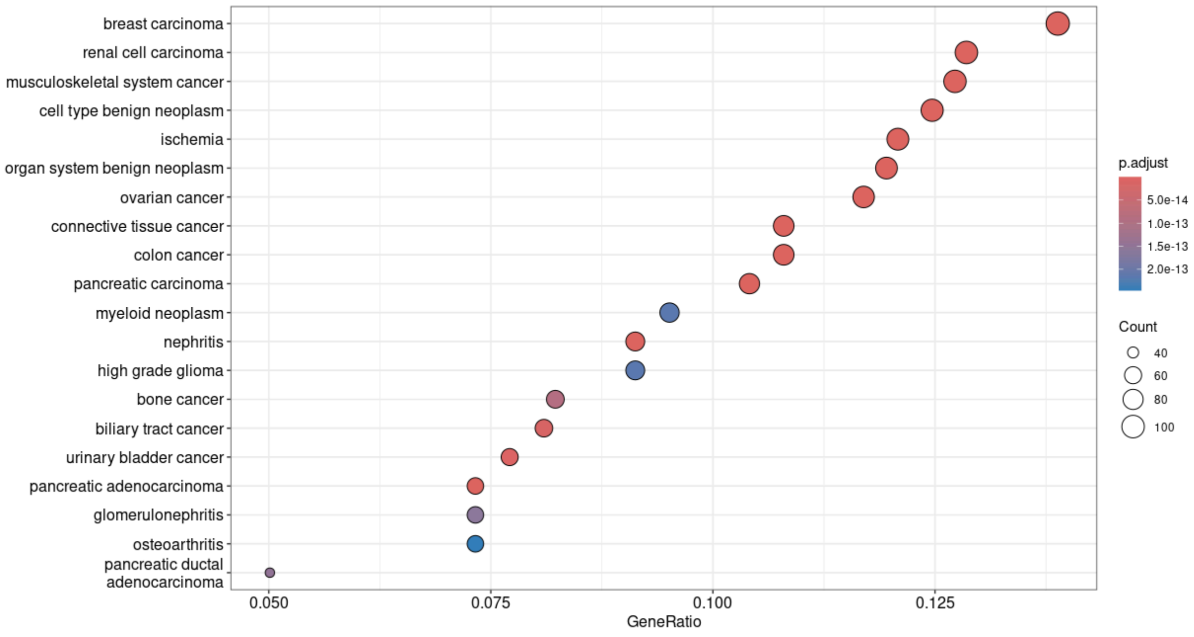


Fig S3. Overrepresentation results with respect to Disease Ontology. Top 20 most enriched terms are presented.

### Tables

| Phylum | Percentage of *UHGPAmyloids* from the phylum | Fisher test p-value | Category |
|---|---|---|---|
| Firmicutes | 70.8% | 1.6e-10 | Overrepresentation |
| Proteobacteria | 24.8% | 5e-41 | Overrepresentation |
| Fusobacteria | 1.7% | 1.7e-4 | Overrepresentation |
| Actinobacteriota | 1.4% | 1.5e-12 | Underrepresentation |
| Bacteroidota | 0.9% | 7.5e-67 | Underrepresentation |
| Campylobacterota | 0.2% | 0.45 | - |
| Myxococcota | 0.1% | 0.2 | - |

Table 1. Fisher tests results for different phyla. For each phylum, the statistical difference in abundance was compared between *UHGPAmyloids* and *UHGP*.

| Uniprot ID | Description | Number of interactions |
|---|---|---|
| Q13287 | N-myc-interactor | 684 |
| P18848 | Cyclic AMP-dependent transcription factor ATF-4 | 460 |
| Q9H6Y7 | E3 ubiquitin-protein ligase RNF167 | 418 |
| P13501 | C-C motif chemokine 5 | 417 |
| P30874 | Somatostatin receptor type 2 | 417 |
| O15551 | Claudin-3 | 417 |
| O14493 | Claudin-4 | 417 |
| P21926 | CD9 antigen | 417 |
| Q9H3Z4 | DnaJ homolog subfamily C member 5 | 417 |
| P10082 | Peptide YY | 417 |

| | | |
|---|---|---|
| P49768 | Presenilin-1 | 417 |
| P04156 | Major prion protein | 417 |
| P02745 | Complement C1q subcomponent subunit A | 417 |
| O00264 | Membrane-associated progesterone receptor component 1 | 417 |
| P37840 | Alpha-synuclein | 417 |
| P67809 | Y-box-binding protein 1 | 417 |
| Q01629 | Interferon-induced transmembrane protein 2 | 417 |
| P13164 | Interferon-induced transmembrane protein 1 | 417 |
| Q99942 | E3 ubiquitin-protein ligase RNF5 | 417 |
| P32241 | Vasoactive intestinal polypeptide receptor 1 | 417 |
| P08172 | Muscarinic acetylcholine receptor M2 | 417 |
| P63000 | Ras-related C3 botulinum toxin substrate 1 | 416 |
| P61586 | Transforming protein RhoA | 416 |
| P60953 | Cell division control protein 42 homolog | 416 |
| Q9Y328 | Neuronal vesicle trafficking-associated protein 2 | 416 |
| Q9UKJ5 | Cysteine-rich hydrophobic domain-containing protein 2 | 416 |
| P26583 | High mobility group protein B2 | 416 |
| P04899 | Guanine nucleotide-binding protein G(i) subunit alpha-2 | 416 |
| P63096 | Guanine nucleotide-binding protein | 416 |

| | | |
|---|---|---|
| | G(i) subunit alpha-1 | |
| P21453 | Sphingosine 1-phosphate receptor 1 | 416 |

Table 2. Proteins from *HPAIntestine_filtered* representing top 5 numbers of interactors from *UHGPAmyloids_filtered* (31 proteins in total).

| GO/KEGG Term | Fisher Test p-value after BH correction |
|---|---|
| growth factor binding | 0.004 |
| transmembrane receptor protein kinase activity | 0.003 |
| integrin binding | 0.016 |
| protein tyrosine kinase activity | 0.002 |
| G protein-coupled receptor binding | 0.02 |
| transmembrane receptor protein tyrosine kinase activity | 0.006 |
| protease binding | 0.004 |
| cytokine binding | 0.035 |
| PDZ domain binding | 0.06 |
| transmembrane transporter binding | 0.021 |
| amyloid-beta binding | 0.016 |
| signaling adaptor activity | 0.009 |
| collagen binding | 0.06 |
| GDP binding | 0.016 |
| virus receptor activity | 0.068 |
| exogenous protein binding | 0.068 |
| ephrin receptor binding | 0.029 |
| G protein activity | 0.013 |
| calmodulin binding | 0.019 |
| cell-cell junction organization | 0.002 |

| | |
|---|---|
| receptor-mediated endocytosis | 0.002 |
| cell-matrix adhesion | 0.002 |
| regulation of cell-substrate adhesion | 0.002 |
| cell-cell junction assembly | 0.006 |
| cell-substrate junction assembly | 0.002 |
| cell-substrate junction organization | 0.002 |
| regulation of protein localization to membrane | 0.005 |
| regulation of phosphatidylinositol 3-kinase/protein kinase B signal transduction | 0.002 |
| focal adhesion assembly | 0.003 |
| positive regulation of phosphatidylinositol 3-kinase/protein kinase B signal transduction | 0.002 |
| regulation of protein localization to cell periphery | 0.004 |
| establishment or maintenance of cell polarity | 0.013 |
| substrate adhesion-dependent cell spreading | 0.007 |
| vesicle-mediated transport in synapse | 0.004 |
| integrin-mediated signaling pathway | 0.002 |
| regulated exocytosis | 0.008 |
| regulation of exocytosis | 0.004 |
| regulation of cell-matrix adhesion | 0.006 |
| regulation of epithelial cell migration | 0.002 |
| Proteoglycans in cancer | 0.002 |
| Focal adhesion | 0.004 |
| Rap1 signaling pathway | 0.005 |
| Axon guidance | 0.007 |
| Regulation of actin cytoskeleton | 0.003 |

| | |
|---|---|
| Ras signaling pathway | 0.006 |
| Adherens junction | 0.013 |
| Endocytosis | 0.005 |
| MAPK signaling pathway | 0.003 |
| Leukocyte transendothelial migration | 0.02 |
| Tight junction | 0.053 |
| Phospholipase D signaling pathway | 0.06 |
| Chemokine signaling pathway | 0.051 |
| Fc gamma R-mediated phagocytosis | 0.028 |
| EGFR tyrosine kinase inhibitor resistance | 0.018 |
| Human papillomavirus infection | 0.016 |
| Relaxin signaling pathway | 0.038 |
| Platelet activation | 0.104 |
| Human cytomegalovirus infection | 0.013 |
| Fluid shear stress and atherosclerosis | 0.01 |

Table 3. Fisher tests results for each GO and KEGG term with Benjamini-Hochberg correction (number of tests = 60) with respect to intestinal proteins only (*HPAIntestine_filtered)*.
